# Elevated soluble Galectin-3 as a marker of chemotherapy efficacy in Breast cancer patients; a prospective study

**DOI:** 10.1101/2020.02.27.968073

**Authors:** Arooj Shafiq, January Moore, Aliya Suleman, Sabeen Faiz, Omar Farooq, Adnan Arshad, Mohammad Tehseen, Ammarah Zafar, Syed Haider Ali, Nasir Ud Din, Asif Loya, Neelam Siddiqui, Fatima K. Rehman

## Abstract

**Purpose:** Galectin-3 (Gal-3) is a glycan-binding lectin with a debated role in cancer progression due to its various functions and patterns of expression. The current study investigates the relationship between breast cancer prognosis and secreted Gal-3.

**Methods:** Breast cancer patients with first time cancer diagnosis and no prior treatment (*n*=88) were placed in either adjuvant or neoadjuvant setting based on their treatment modality. Stromal and plasma Gal-3 levels were measured in each patient at the time of diagnosis and then throughout treatment using immunohistochemistry (IHC) and ELISA respectively. Healthy women (>18 years of age, *n*=63) were used to establish baseline levels of plasma Gal-3. Patients were followed for 84 months for disease free survival analysis.

**Results:** Enhanced levels of plasma (adjuvant) and stromal (neo-adjuvant) Gal-3 were found to be markers of chemotherapy efficacy. The patients with chemotherapy induced increase in extracellular Gal-3 had longer disease-free interval and significantly lower rate of recurrence during 84-month follow-up compared to patients with unchanged or decreased secretion.

**Conclusion:** The findings support the use of plasma Gal-3 as a marker for chemotherapy efficacy when no residual tumor is visible through imaging. Furthermore, stromal levels in any remaining tumors post chemotherapy can also be used to predict long term prognosis in patients.

**Key points:** Increased Gal-3 secretion due to chemotherapy leads to better prognosis and longer disease-free survival.

The analysis of soluble Gal-3 expression could be useful as a support tool in predicting treatment efficacy in patients with no visible tumor remaining for follow up through imaging.

## INTRODUCTION

While there have been significant advances in technology to diagnose breast cancer, accurate prognostic tools during treatment remain lacking. Currently, anticancer treatment varies according to cancer type and grade. A low-grade or localized tumor may be treatable with minimal chemotherapy after surgical removal (adjuvant) while a high-grade tumor may require initial aggressive chemotherapy to shrink the mass prior to surgery (neo-adjuvant) [1]. Monitoring of therapy efficacy is essential, so that treatment may be continually optimized to reduce remaining cancer burden and possibility of disease relapse [2,3]. However, continuing presence of cancer has proven difficult to determine. Various measurements have been developed, including 1) size and cellularity of the primary tumor and nodal metastases [3]; 2) imaging techniques such as MRI and PET/CT imaging [2,4]; and 3) circulating tumor cell DNA (ctDNA) detection [5]. Tissue-based biomarkers such as Estrogen receptor (ER), Progesterone receptor (PR) and Human epidermal growth factor receptor (HER2) and measurement of mitotic activity through Ki67 assay are also commonly used at the time of diagnosis to determine effectiveness of treatment options [2,6]. More recently, absence of Galectin-3 (Gal-3), a glycan-binding lectin, has been associated with higher *in vivo* growth in murine breast tumors [7] and poor prognosis in node-positive breast cancers [8].

Gal-3 is a 31 kDa glycoprotein that interacts with targets in both the extracellular and intracellular space to regulate various biological pathways including cell growth, differentiation, adhesion, inflammation and apoptosis. It can be found on the cell surface, in the cytoplasm and nucleus, and is also secreted into the extracellular space [9]. Studies have shown conflicting results regarding Gal-3 function in tumor formation, growth and progression depending on localization [10,11]. For example, elevated expression of intracellular and extracellular Gal-3 has been associated with resistance in thyroid [12,13] and breast [14] cancers, and promotion of metastasis in pancreatic cancer [15,16]. On the other hand, extracellular secreted Gal-3 has been found to induce apoptosis in some cell lines, including human T cell leukemia and peripheral blood mononuclear cells [17]. Furthermore, downregulation of intracellular Gal-3 has been associated with metastasis in breast [7,18] and gastric [19] cancers.

Previous studies have also looked at regulation of Gal-3 expression and localization. One particular study found Gal-3 in the media from cancer cells after p53 activation [20], indicating possible role in cell cycle regulation or apoptosis. However, Gal-3 is not a transcriptional target of p53 and overall expression remain similar regardless of p53 status. Instead, it is secreted through a non-classical exosome-mediated pathway largely controlled by TSAP6 protein, which is a transcriptional target of p53 [21].

Many chemotherapy drugs work more effectively in tumors with active p53 pathway. Since extracellular Gal-3 has been implicated in apoptosis we speculated if it may play a role in tumor response to chemotherapy. Here, we investigate soluble Gal-3 as a possible biomarker for chemoefficacy in breast cancer patients undergoing chemotherapy and surgery in the adjuvant or neo-adjuvant setting. Based on the current literature, we hypothesize an increase in soluble Gal-3 levels in tumor stroma or plasma of chemoresponsive patients.

## MATERIALS AND METHODS

### Sample Collection and Patient Information

This study protocol was approved by the Shaukat Khanum Memorial Cancer Hospital and Research Center Institutional Review Board (IRB protocol number 07-01). Breast cancer patients with first time diagnosis of cancer and no prior treatment were recruited with informed consent to this two-site, prospective randomized clinical study between March 2007 and March 2008. Sample collection continued until all required samples over the course of their therapy were collected or a patient withdrew from the study. Eighty-eight (88) adult female patients presenting to Shaukat Khanum Memorial Cancer Hospital and Research Center (SKMCH&RC) were included in the final study. Patients included in the study received a combination of chemotherapy and surgical tumor removal. All patients in the study received taxane therapy as part of their treatment protocol. Blood samples were collected from all 88 patients at the time of diagnosis prior to therapy. Non-tumoral control amples (*n*=63) were obtained from healthy women at SKMCH&RC (*n*=32) and the Institute of Biochemistry and Biotechnology, University of the Punjab, Lahore, Pakistan (*n*=31).

Cancer patients were divided into adjuvant (n=52) and neo-adjuvant setting (n=36) based on their treatment protocol. Blood samples were taken in both groups at each chemotherapy visit as well as immediately prior to surgery and again two weeks post-surgery. Blood specimens collected in EDTA-coated tubes were immediately centrifuged at 2000 x g for 10 minutes and separated plasma was stored at −70°C until use. Tumor and adjacent normal breast tissue were collected during surgical removal. Part of the sample was frozen and stored in liquid nitrogen while the remaining was paraffin-embedded for future use. Summary of patients in each group is listed in Table 1. Details of patients in each group including age at disease onset, tumor size and histologic type, lymph node involvement and ER/PR and Her2 status, are listed in Supplemental Tables I and II. Patients were followed for 84 months following end of treatment for disease free survival analysis. Survival time was defined as the period of time in months from the date of diagnosis to the date of death from breast cancer. Relapse time was defined as the period of time in months from the time of diagnosis to the date at which relapse was clinically identified. All efforts were made to follow REMARK guidelines during study design and analysis [22].

**Table I:** Summary of patient characteristics for breast cancer patients enrolled in the study. Patient age, tumor size, type, grade and metastatic status are given. TRN=Triple negative; ER= Estrogen receptor; PR= Progesterone receptor; IDC=Intra-ductal carcinoma; DCIS= Ductal carcinoma in-situ; LCA= Lobular carcinoma.

|  | **Adjuvant**<br>**n=52** | **Neo-Adjuvant**<br>**n=36** |
| --- | --- | --- |
| **Age (years)** |  |  |
| **Mean** | 45.8 | 44.1 |
| **Median** | 45 | 45.8 |
| **Range** | 28-65 | 20-62 |
| **Size (cm<sup>3</sup>)** |  |  |
| **Mean** | 23.3 | 34.2 |
| **Median** | 19.3 | 15.9 |
| **Range** | 0.24-106.2 | 1.1-225 |
| **Axillary nodes n(%)** |  |  |
| **Yes** | 36 | 18 |
| **No** | 14 | 14 |
| **Unknown** | 2 | 4 |
| **Genetic background (n)** |  |  |
| **TRN** | 9 | 11 |
| **Her2+** | 6 | 8 |
| **ER+** | 20 | 11 |
| **HER2+/ER+** | 10 | 1 |
| **Unknown** | 7 | 5 |
| TRN=Triple negative; ER= Estrogen receptor; PR= Progesterone receptor; IDC=Intra-ductal carcinoma; DCIS= Ductal carcinoma in-situ; LCA= Lobular carcinoma. |  |  |

### Protein quantification in Crude Plasma Samples

Crude plasma protein concentration was calculated by Bradford assay [23] as per manufacturer’s protocol. Briefly, 5 µl of each sample was added to 1 ml of Bradford reagent. Samples were incubated for 10 minutes before measuring absorbance at 595 nm on BioRad-Smart Spec plus spectrophotometer (BioRad, USA). Protein concentration was calculated against a standard curve drawn using serial dilutions of BioRad BSA standard solution.

### Immunohistochemical analysis (IHC)

Paraffin-embedded tissue samples from all study patients were examined using hematoxylin and eosin staining to confirm tumor grade and type. Only one sample at the time of surgery was obtained for patients in adjuvant setting while two samples were taken from neo-adjuvant patients; one at the time of diagnosis (Pre) and a second at the time of surgery after chemotherapy (Post). Gal-3 and Ki67 expression was detected by IHC using mouse polyclonal anti-Gal-3 (Santa Cruz, CA, USA, SC-14364, 1:500) and rabbit polyclonal anti-Ki67 (abcam, Cambridge, MA, USA, ab92742, 1:2500) antibodies respectively. After standard deparaffinization, samples were tested using Envision dual link dakocytomation kit following the manufacturer’s protocol (Dako-Behring, Glostrup, Denmark). An irrelevant IgG was used as the primary antibody in controls. Scoring was performed by three blinded observers; two researchers and one pathologist in 3 different fields (at 200x and 400x magnification) for each specimen. Mean score by the three observers was recorded as the final result.

### Galectin-3 ELISA

Secreted Gal-3 concentrations in plasma samples were quantified by ELISA (Bender MedSystems, Austria) using provided manufacturer protocol. Briefly, various dilutions and undiluted plasma from each patient or control sample was incubated for 2 hours with biotin-conjugated Gal-3 antibody followed by addition of streptavidin-HRP. Standard curve to extrapolate actual amount in samples was drawn using serial dilutions of provided recombinant Gal-3. All samples were tested in triplicate. The ELISA used incorporated TMB substrate to reach a chromagenic end point detected at 450 nm (reference wavelength = 620nm) on BioRad microplate reader (Microplate manager 5.2.1).

### TUNEL staining for cell death

Apoptosis in frozen tissue sections was measured by TUNEL staining using “*In Situ* Cell Death Detection Kit, AP” (Roche, Germany). After target retrieval, high (70W) microwave irradiation was applied for 1 minute. The samples were then blocked in 0.1M Tris-HCl (pH: 7.5), containing 3% BSA (w/v) and 20% (v/v) normal bovine serum. Slides were rinsed with 1X PBS, blocked and stained with TUNEL mixture. The fluorescent stain was converted to chromagen based results using AP-converter. Finally, slides were counter stained with 0.1% eosin (v/v), mounted with aqueous mounting media and analyzed under light microscope.

### Statistical analyses

Plasma and stromal Gal-3 was measured as a continuous score and analysis was undertaken by log transforming Gal-3 and using this log(Gal-3) as a covariate to investigate whether there is a linear increase in the probability of relapse with increasing Gal-3 value. Statistical analyses were carried out using Statview and Excel data analysis package. As there was no clinically defined cutoff point for plasma Gal-3 level, values above 95% of the median non-tumoral control values were considered elevated. In addition, change in Gal-3 plasma levels in response to taxane therapy was used to divide the patients into two groups (increased versus unchanged/decreased serum Gal-3 levels). The influence of plasma Gal-3 levels on eventual recurrence was tested using a simple binary indicator as the variable which was positive only when the levels showed at least a two-fold increase over time. The resulting contingency table was analyzed using Fischer’s exact test. Next, we assessed if the mean stromal levels of Gal-3 varied significantly between patients with or without recurrence using Levene’s and Kolmogorov-Smirnov tests to check for equal variance and normality assumption respectively. Two-tailed unpaired T-test was used to compare the statistical significance of the differences in data from two groups, where appropriate using Excel statistics package. A Cox regression model was used with soluble Gal-3 as the exposure variables and overall survival (from time of surgery to time of death or end of current follow-up) as the outcome. Disease-free survival (DFS) was defined as the time from surgery to the first one of the following events: recurrence at local or distant sites, or death without recurrence. Patients were excluded from the survival analysis if they were lost to follow up. Survival curves were plotted by the Kaplan–Meier method and compared using the log-rank test in SPSS software (SPSS Inc., Chicago, IL, USA); p < 0.05 was considered as statistically significant. The analyses were adjusted simultaneously for sex, age, tumor size (World Health Organization), and stage. Chemotherapy induced elevated levels of Gal-3 were statistically analyzed through single factor ANOVA and paired t-tests. Pearson correlation coefficient was used to measure relationship between Gal-3 levels and apoptosis measured through TUNEL reactivity pre vs. post-chemotherapy.

## RESULTS

### Extracellular Gal-3 levels decrease with increasing grade of breast cancer

Several studies have examined the expression of Gal-3 in tumor samples from multiple tissues. Most find levels of intracellular Gal-3 increased with metastasis and advanced cancer [12-16]. However, very few have examined levels of extracellular Gal-3. Therefore, we first examined the levels and expression pattern of Gal-3 in study patients’ tumor biopsies collected at the time of diagnosis (n=88). Gal-3 was typically absent in stroma of normal breast tissue. In contrast, significant levels of extracellular Gal-3 were observed in the stroma of low-grade (Grade I & II) breast cancer cells (Fig 1a & b). These levels decreased dramatically with increasing malignancy grade (*p*<0.05) (Fig. 1b). These findings are consistent with results in other studies involving breast [8,18] and endometrial cancers [24].

**Figure 1:**
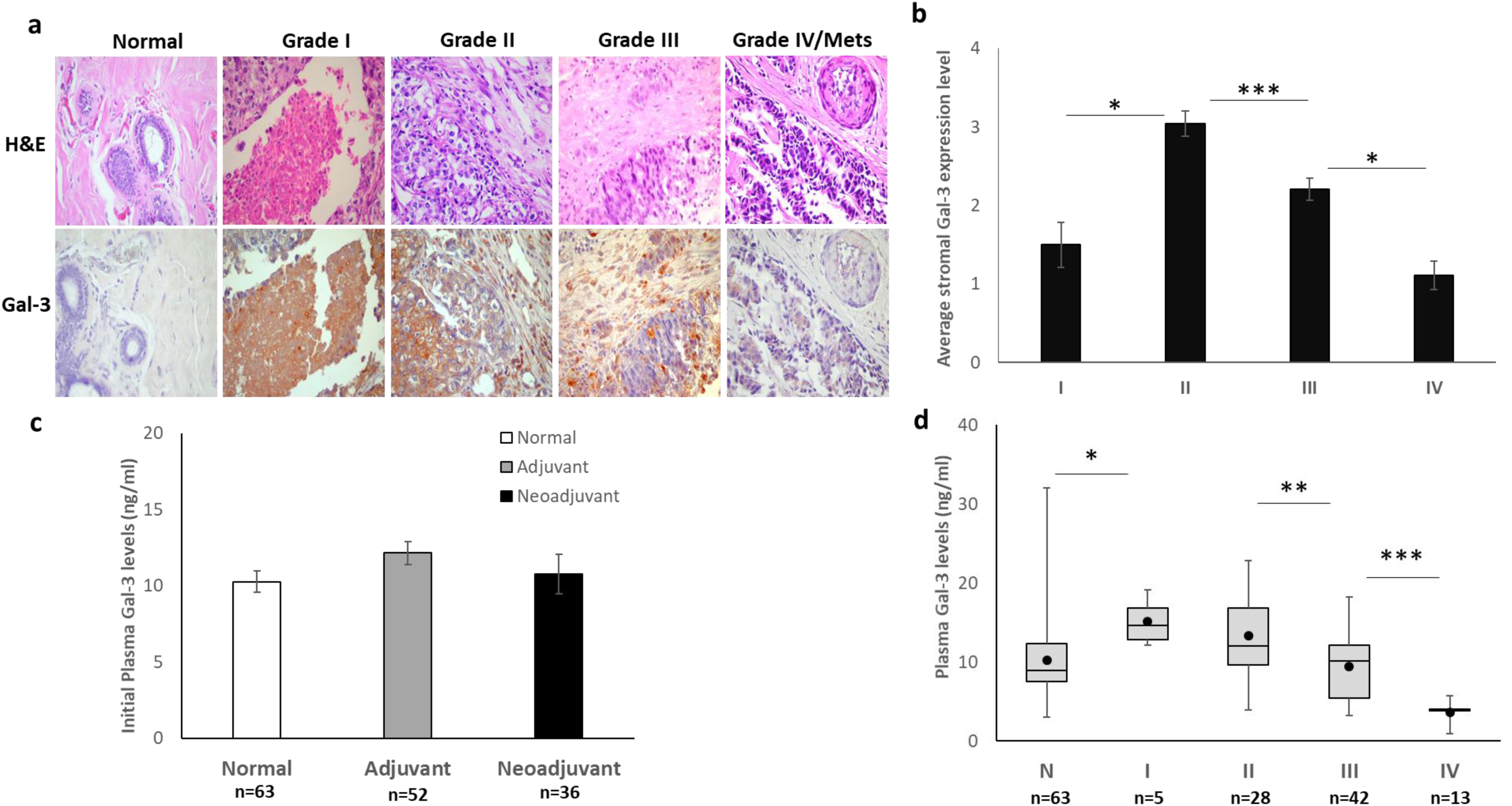
Stromal Galectin-3 (Gal-3) levels decrease with increasing tumor grade in breast cancer patients: **a.** Gal-3 localization and expression levels were examined through immunohistochemistry (IHC) of paraffin-embedded tissue sections from all study patients (n=88) at the time of diagnosis prior to any treatment. Normal breast tissue has undetectable levels of Gal-3 while low grade tumors show high Gal-3 expression in stroma and extracellular spaces. These levels decline with increasing grade. Representative images from each category are shown at 400x final magnification. **b.** Average stromal Gal-3 expression levels are shown in Grade I-IV breast carcinoma tissues from our study patients (n=88). Gal-3 levels in stroma were quantified as 0-4 based on expression intensity. Average of three separate blinded observations is shown. Expression levels rose between Grade I and II (p<0.01) and then dropped significantly with increasing grade (p<0.0001 Grade II vs. III; p<0.01 Grade III vs. IV). **c.** Gal-3 concentration in plasma from study patients was quantified through ELISA in the various study groups. The average levels remained unchanged prior to therapy compared to non-tumoral controls. Each plasma sample was quantified in triplicate (n=63 nontumoral controls; n=52 Adjuvant; n=36 Neo-adjuvant patients). **d.** Average Gal-3 plasma levels in each category are negatively correlated to tumor grade. Plasma levels of Gal-3 were quantified by ELISA. Gal-3 levels were found to be significantly elevated in plasma from low grade breast cancer patients (Stage I & II (n=33)) prior to any treatment compared to nontumoral controls (n=63). The Gal-3 levels dropped sharply in Grade IV and metastatic cancers (n=11; p<.05). Low levels of Gal-3 (mean = 10.6 ng/ml; range: 3.0-34.0 ng/ ml) were observed in healthy adult women (n=63) plasma. ng/ ml (95^th^ percentile) was considered to be the top limit of normal levels in our analysis. Error bars show +/-SEM.

Next, we examined if differential Gal-3 levels could be observed in the plasma of patients as well. Total protein was first quantified from the plasma of breast cancer patients (n=88) and from healthy female controls (n=63). Total plasma protein levels remained similar in all study groups (Supplemental fig. 1a). Similarly, the initial plasma Gal-3 levels remained unchanged in both adjuvant (n=52) and neo-adjuvant (n=36) groups when compared to non-tumoral controls (n=63) (Fig. 1c) ELISA results showed a median of 11.0 ng/ml Gal-3 protein in 90% of the control women in accordance with previous findings [25]. There was a robust increase in plasma Gal-3 levels in low grade (grade I & II) tumors compared to non-tumoral controls. Similar to observations from tumor stroma, the levels of plasma Gal-3 had an inverse relationship with increasing grade of breast cancer (Fig. 1d). These results were confirmed by western blot analysis (Supplemental fig. 1b).

### Elevated Gal-3 levels are predictive of patient response when receiving adjuvant chemotherapy

As levels of secreted Gal-3 in stroma and plasma of breast cancer patients were altered with varying grades, we next examined if stromal Gal-3 levels change with chemotherapy and if they could be correlated to response in patients receiving adjuvant treatment following surgical removal of primary tumor. Tumor samples were taken before any chemotherapy was administered and stained with H&E and Gal-3 to visualize expression pattern and localization of the Gal-3 protein. High Gal-3 levels were seen in the tumor stroma of 45% of these patients while the remaining samples either had no expression or exhibited mostly intracellular Gal-3 with minimal release in the extracellular space (Fig. 2a). Patients that remained in remission over the 84 month follow up period were more likely to have moderate to high levels of stromal Gal-3 in the primary tumor (Fig. 2b; p=.0009).

**Figure 2:**
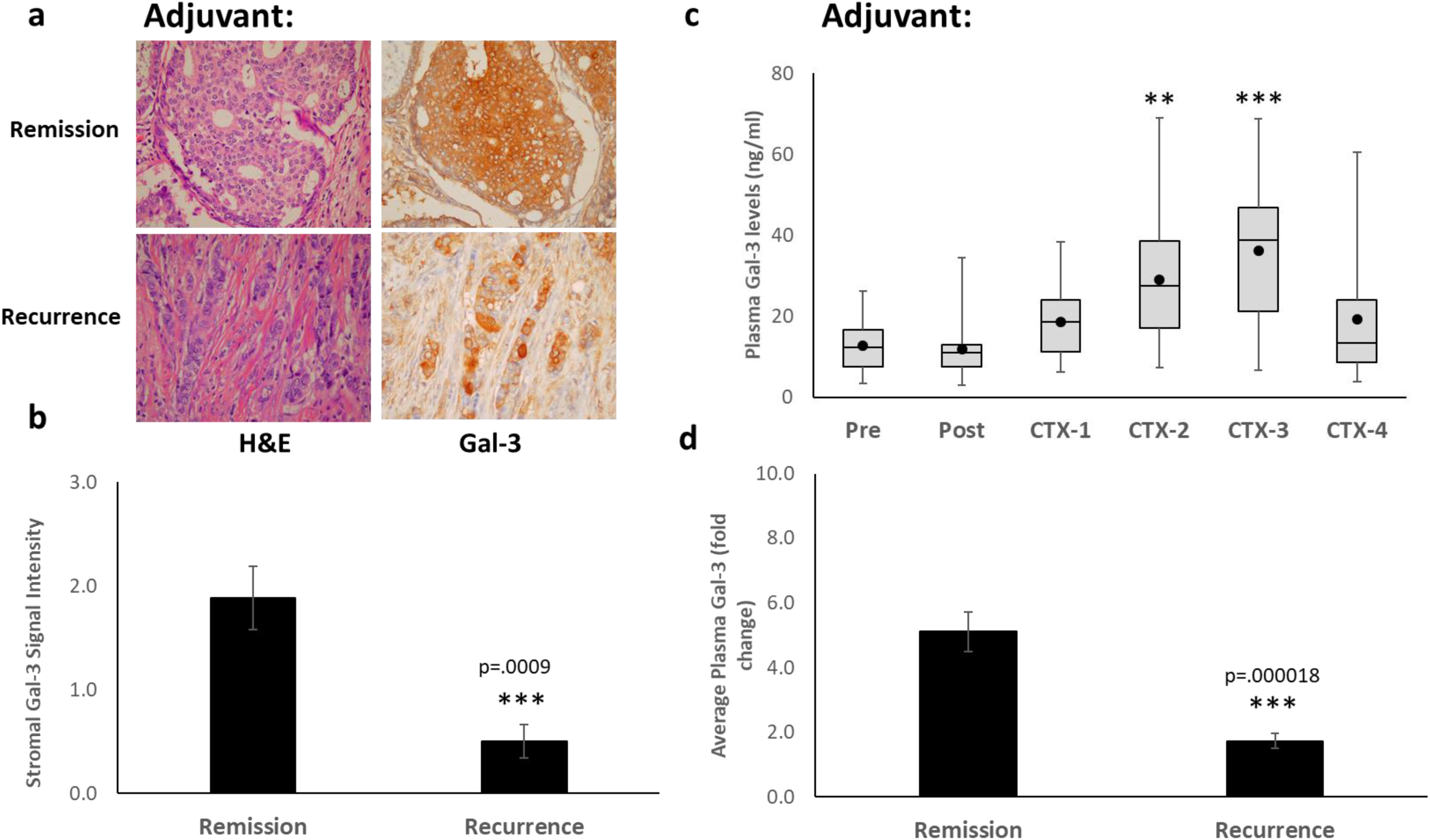
Elevated levels of Gal-3 are a positive predictor for response for patients receiving adjuvant chemotherapy: **a&b** Sections of paraffin-embedded tumor tissue were stained with H&E for general morphology and Gal-3 for expression level and localization analysis (400x magnification). Each stained sectioned was scored by two researchers and a pathologist. Patients remaining in remission at the end of 84 month follow up period had significantly higher levels of extracellular Gal-3 in tumor microenvironment compared to those with recurrent/ metastatic disease (p=.001). **c.** Average plasma Gal-3 levels were calculated at the time of diagnosis (pre), 2 weeks post-surgical tumor removal (Post) and after each chemotherapy cycle (CTX-1-4) in patients receiving adjuvant chemotherapy (n=45). Notice significant increase in Gal-3 levels with each chemotherapy cycle in adjuvant patients (Pre:CTX2 p=0.008812; Pre:CTX3 p=0.000717 compared to initial (Pre) levels). **d.** Patients that went in remission and remained disease free over the 84 month follow up period had at least a 2-fold increase (range 2-11 fold) in plasma Gal-3 levels in response to chemotherapy. On the other hand, patients with recurrent disease did not show the same dramatic increase in plasma Gal-3 levels post chemotherapy (p=.000018). Error bars show +/-SEM.

Next, plasma samples were taken at the time of diagnosis, two weeks post-surgery and after each chemotherapy session in order to examine if plasma Gal-3 levels were also altered in response to chemotherapy. Gal-3 levels in plasma were found similar to non-tumoral control prior to chemotherapy but rose significantly once treatment started (p=0.009 and p=0.007 after chemotherapy cycles 2 and 3 (CTX-2 and CTX-3) respectively compared to initial (Pre) levels (Fig. 2c & Table 2; Supplemental Fig. 2b). The ELISA results were confirmed by western analysis (Supplemental Fig. 2a). Marked increase in plasma Gal-3 levels was measured in 59.6% (31/52) of the patients while the remaining had no evident change in response to chemotherapy. Patients remaining disease free at 84 months presented with a 5-fold elevated plasma Gal-3 on average (range 2.5 - 11.0 fold) compared to their pre-chemotherapy levels (Fig. 2d; p=.000018). These results imply that plasma levels of Gal-3 may be a useful predictor of chemotherapy efficacy especially in cases where no residual tumor is left to follow through imaging.

**Table II:**
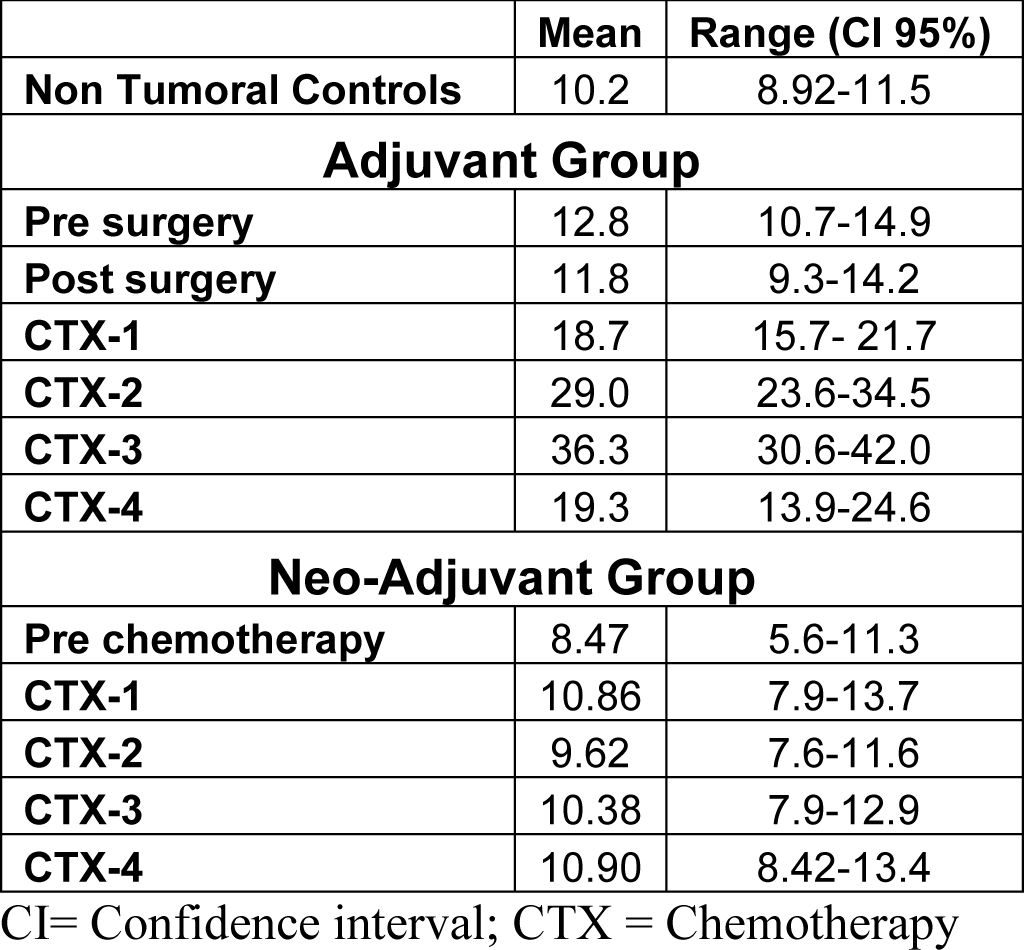
Plasma Gal-3 level distribution in study population. CI=Confidence interval; CTX = Chemotherapy.

| **Plasma Gal-3 Levels in Study Population** |  |  |
| --- | --- | --- |
|  | **Mean** | **Range (CI 95%)** |
| **Non Tumoral Controls** | 10.2 | 8.92-11.5 |
| **Adjuvant Group** |  |  |
| **Pre surgery** | 12.8 | 10.7-14.9 |
| **Post surgery** | 11.8 | 9.3-14.2 |
| **CTX-1** | 18.7 | 15.7- 21.7 |
| **CTX-2** | 29.0 | 23.6-34.5 |
| **CTX-3** | 36.3 | 30.6-42.0 |
| **CTX-4** | 19.3 | 13.9-24.6 |
| **Neo-Adjuvant Group** |  |  |
| **Pre chemotherapy** | 8.47 | 5.6-11.3 |
| **CTX-1** | 10.86 | 7.9-13.7 |
| **CTX-2** | 9.62 | 7.6-11.6 |
| **CTX-3** | 10.38 | 7.9-12.9 |
| **CTX-4** | 10.90 | 8.42-13.4 |

CI= Confidence interval; CTX = Chemotherapy

### Gal-3 release in tumor stroma can be used to monitor chemotherapy efficacy in Neo-adjuvant setting

In order to elucidate if the elevated plasma and stromal Gal-3 were enhanced by chemotherapy itself, we next measured the Gal-3 levels in patients receiving taxane therapy prior to primary tumor resection (Neo-adjuvant setting). Gal-3 levels and expression pattern in tumor tissue were assessed in initial diagnostic biopsy (pre-chemotherapy) and in the surgically removed tumor post-chemotherapy. Generally, low levels of stromal Gal-3 were observed in almost all patients pre-chemotherapy (n=36) (Fig. 3a & b top panels). During the 84-month follow up period, patients remaining in remission had a 2-fold or more increase in stromal Gal-3 levels in response to chemotherapy (Fig. 3a bottom panel & 3c, p=.030). On the other hand, patients with recurrent disease typically showed no change in Gal-3 levels or expression pattern with chemotherapy administration (Fig. 3b bottom panel & 3c). This data indicates stromal Gal-3 as a positive marker of chemotherapy response in neo-adjuvant patients as well. However, unlike adjuvant setting, no enhancement or significant change in plasma levels of Gal-3 were observed in patients receiving neo-adjuvant chemotherapy (Fig. 3d & Supplemental Fig. 2c).

**Figure 3:**
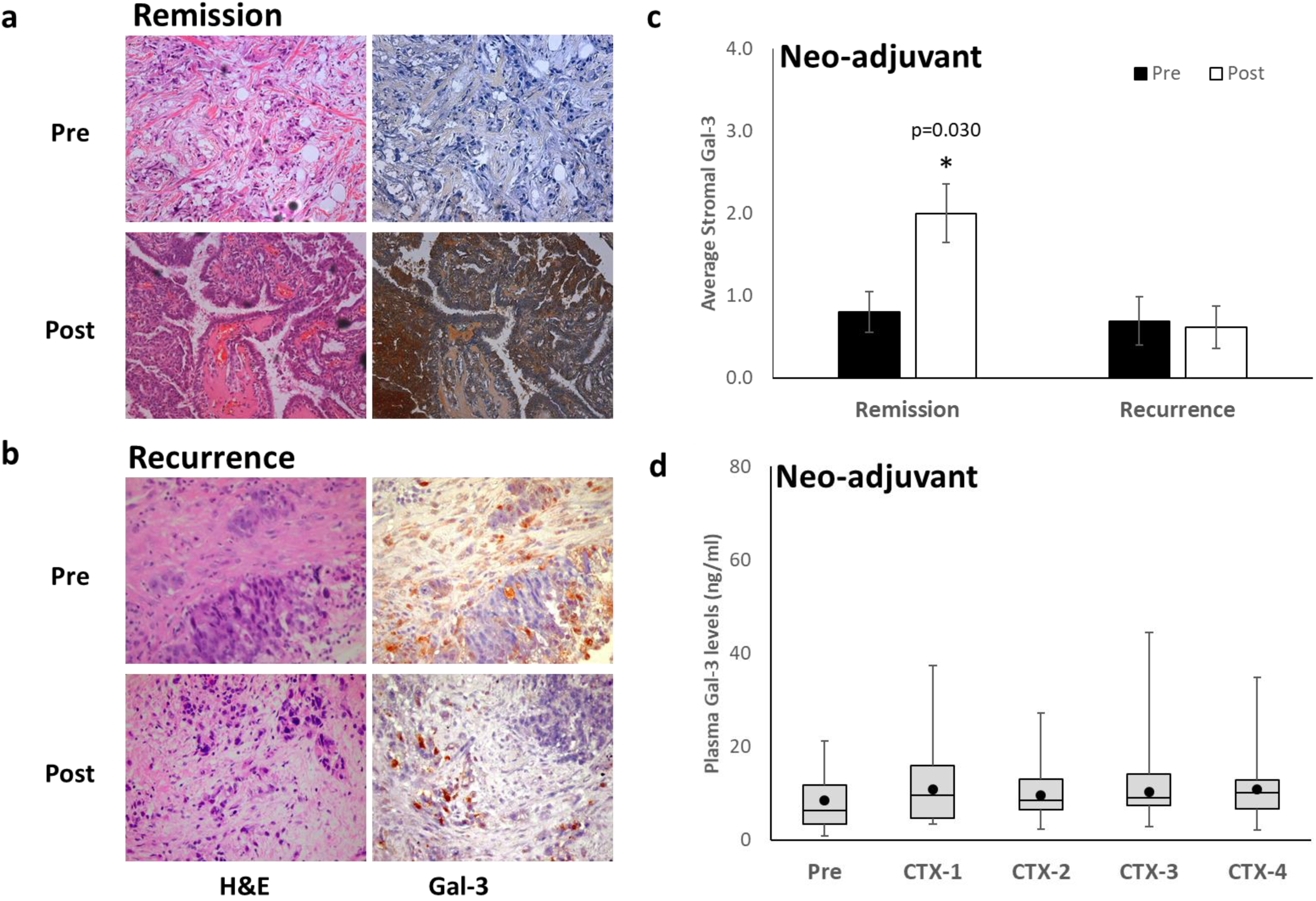
Elevated extracellular Gal-3 is a marker of chemotherapy efficacy in breast cancer patients receiving neo-adjuvant chemotherapy: Levels of extracellular Gal-3 were monitored in patients receiving chemotherapy prior to surgery before (pre) and after (post) chemotherapy through immunohistochemical (IHC) analyses on paraffin-embedded biopsy (pre-chemo) and surgical (post-chemo) samples. **a&c.** Most of the patients that remained in remission during the study period exhibited high expression of Gal-3 in tumor stroma after taxol therapy. **b&c.** This increase in extracellular Gal-3 levels was not observed in patients with disease recurrence within 20 months. **c.** This inhibition of cancer recurrence was true regardless of the initial grouping and was only dependent upon the increase in stromal levels of Gal-3 post Taxane chemotherapy (p=.030). **d.** Average plasma Gal-3 levels remained low and unchanged in response to chemotherapy.

### Extracellular Gal-3 in tumor stroma correlates with apoptosis in response to chemotherapy

To determine if augmented levels of stromal Gal-3 correlate with higher rate of apoptosis in tumor post chemotherapy, tissue sections were examined with TUNEL assay. Results were quantified as the percentage of tumor cells with TUNEL positive nuclei. Higher levels of TUNEL staining were seen in patients with increased levels of stromal Gal-3 post chemotherapy, indicating improved apoptosis (Supplemental Fig. 3a & b). Pearson’s correlation tests also show positive correlation between stromal Gal-3 levels and TUNEL reactivity in post chemotherapy samples (Supplemental fig. 3c & d; pearson coefficient correlation = 0.730 post chemotherapy). These results propose a possible role of extracellular Gal-3 in induction of tumor apoptosis in response to chemotherapy.

### Extracellular secreted Gal-3 in response to Taxane therapy is prognostic of disease-free survival

Finally, we assessed the prognostic value of the elevated Gal-3 levels in plasma (adjuvant; n=45) and tumor stroma (neo-adjuvant; n=31) through Kaplan-Meier survival analysis after 84-month follow up of study patients (Fig. 4a & b; Fig. 5). In the adjuvant setting, patients with increased plasma levels of Gal-3 in response to chemotherapy had a significantly longer disease-free interval (22 months versus 11 months) as well as disease-free survival (77.4% versus 50.0%) compared to post-chemotherapy patients with decreased or unchanged Gal-3 levels (Fig. 4a; p<0.01). Neoadjuvant study group patients had an even longer disease-free interval (35 months versus 6 months) and higher disease-free survival (81% versus 40%) (Fig. 4b; p<.0001) in the presence of elevated Gal-3 in tumor stroma. These results help support presence of extracellular Gal-3 as a positive indicator of prognosis.

**Figure 4:**
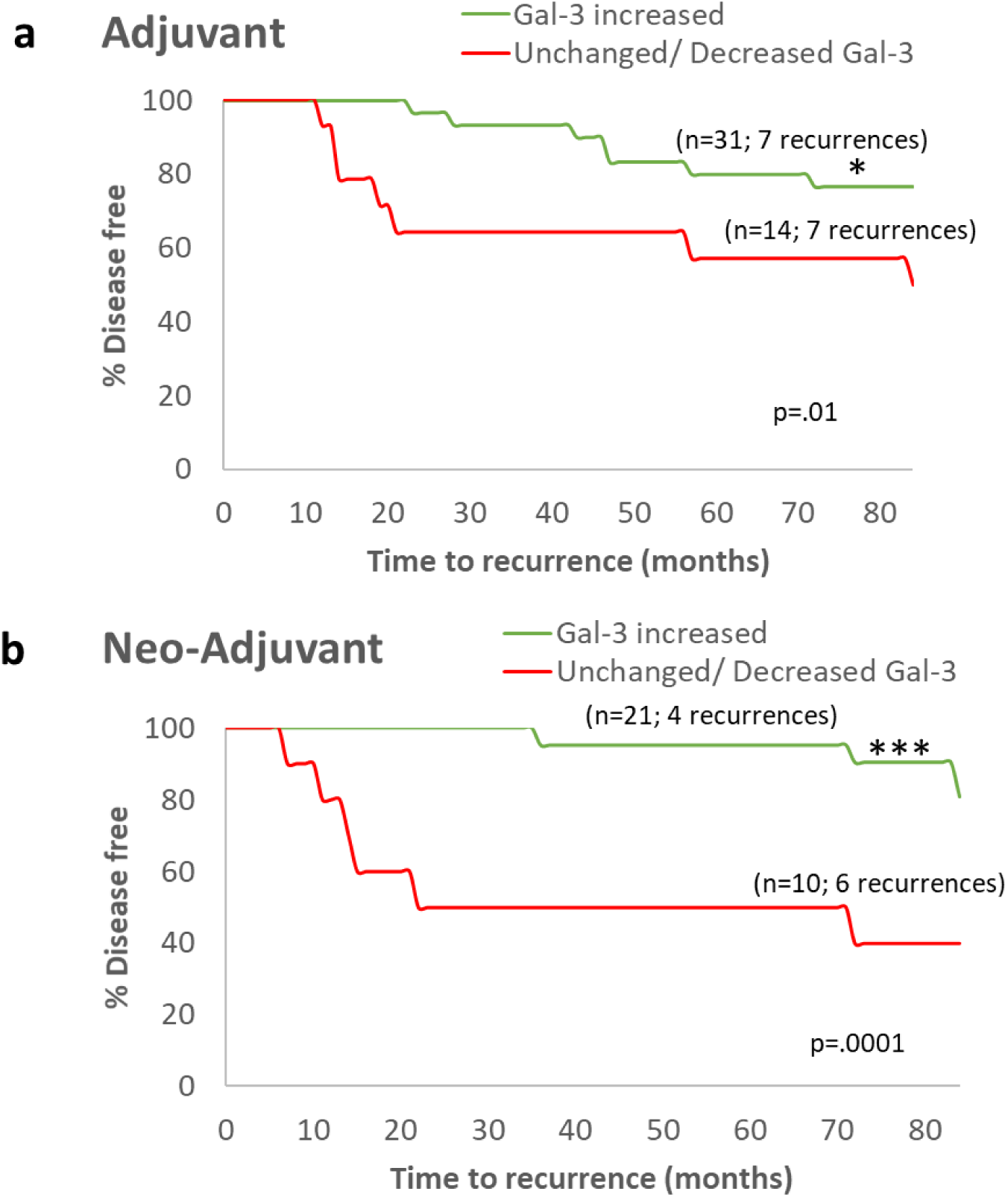
Taxane therapy induced Gal-3 secretion is prognostic of disease-free survival (DFS): Patients with increased Gal-3 expression in both Adjuvant **(a)** and Neo-adjuvant **(b)** settings showed increased DFS over the study period of 84 months. These patients additionally showed a longer interval without recurrence compared to patients with unchanged or decreased Gal-3 (p<0.01 (adjuvant); p<0.0001 (neoadjuvant)).

**Figure 5:**
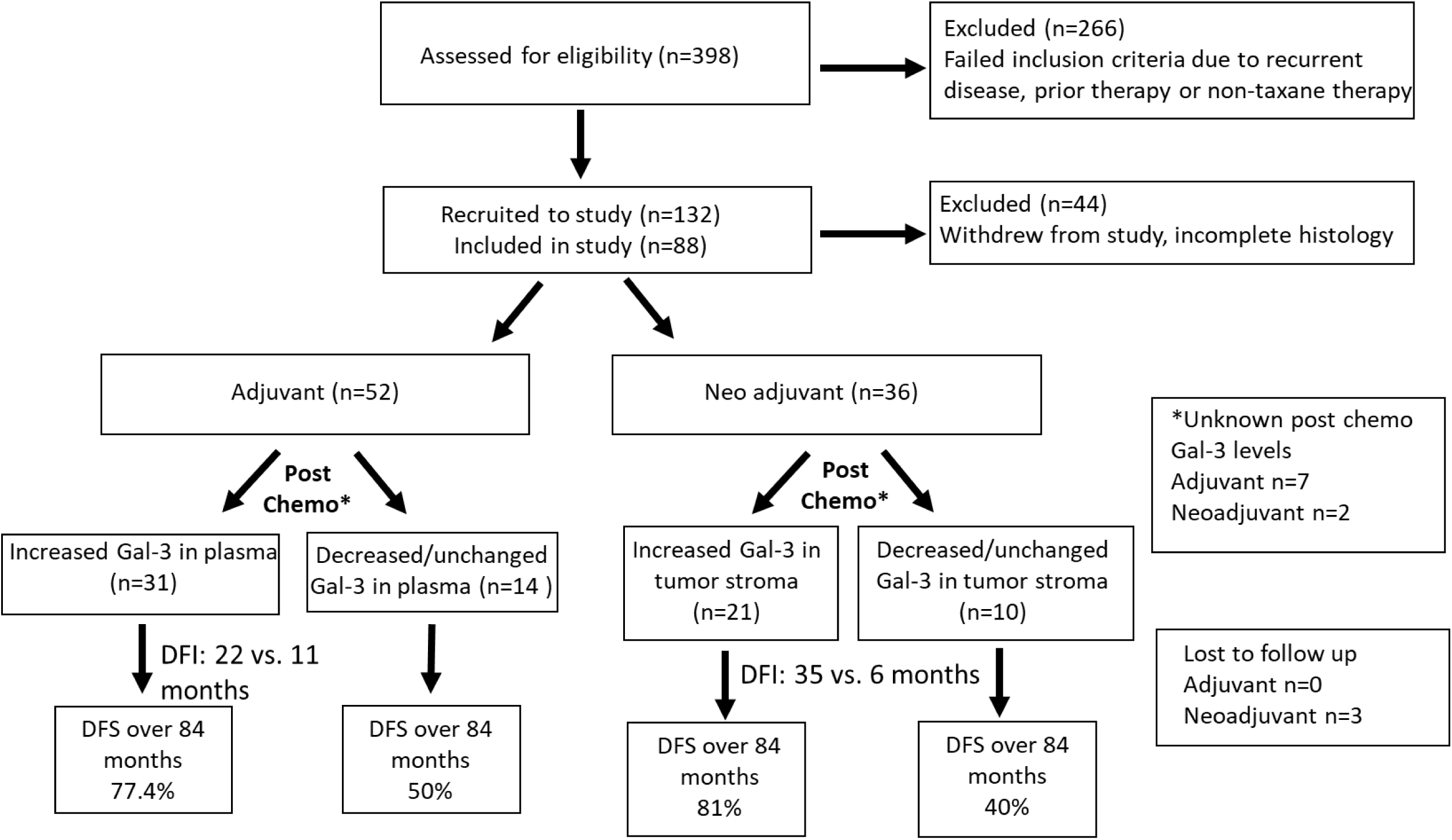
Flow chart of study progress, including recruitment and monitoring of study participants: Study participants consisted of non-tumoral controls (n=63) and patients with first time breast cancer diagnoses (BCD) (n=88). Patients with BCD were further divided into two group based on treatment modality (Adjuvant chemotherapy, n=52 vs. Neo-adjuvant chemotherapy, n=36). All patients in the study received taxane therapy as part of their chemotherapy regimen. Stromal and plasma levels of Gal-3 were measured at the time of recruitment and throughout the treatment period. Following end of treatment, patients were monitored for a total of 84 months for disease-free survival (DFS).

## DISCUSSION

In the present study, we analyzed stromal and plasma Gal-3 levels in response to adjuvant and neo-adjuvant treatment in newly diagnosed breast cancer patients. We hypothesized that changes in plasma Gal-3 levels over the course of treatment may correlate with disease prognosis and treatment efficacy. Analysis of the tumor tissue prior to treatment showed a clear inverse relationship between levels of secreted Gal-3 in untreated tumor tissue and stage of malignancy. This result was in agreement with a recent study by Ilmer et al. (2016), which demonstrated a trend of decreased Gal-3 in higher-grade breast tumors, and significant association of decreased Gal-3 with lymphovascular invasion [8]. Similar results have also been seen previously by Castronova et al. in breast cancer and Okada et al. in gastric cancers [18,19].

In contrast to previous published studies, we followed the expression and localization of Gal-3 over the course of treatment in our study patients. In the neo-adjuvant setting, significant increase of stromal Gal-3 was seen in nearly 60% of the study population post-chemotherapy (Fig. 5). The higher concentration may signify that chemotherapy induced secretion of Gal-3 in this group. However, plasma levels remained unchanged, implying that secreted Gal-3 may bind to partners within the tumor microenvironment or the tumor cells themselves. Few comparative studies have been found that distinguish between stromal and plasma Gal-3 levels in the settings of neo-adjuvant versus adjuvant taxane treatment, especially in patients with no prior treatment history. Some studies specified parameters within which Gal-3 was examined, such as *in vitro, in vivo*, post-chemotherapy [8] and no chemotherapy [7,19]. Many of the studies however used more relaxed parameters for increased study participation, thus resulting in limited insight into changes in Gal-3 localization and function in response to treatment. The study by Ilmer et al. used very specific inclusion criteria (no prior malignant history and doxorubicin therapy) and found increased tumorigenicity and drug resistance in patients exhibiting loss of Gal-3, leading to poor prognosis [8]. These results were comparable to the present study findings, where an inverse relationship between levels of soluble Gal-3 and disease progression is demonstrated. However, the Ilmer et al. study was a retrospective analysis, and pre-therapy or changing levels in the plasma were not examined.

In order to elucidate the exact effect of chemotherapy on Gal-3 expression and localization, plasma levels of Gal-3 were examined post each chemotherapy cycle. Plasma levels of Gal-3 did not change significantly compared to non-tumoral controls in neo-adjuvant patients. The latter observation suggests that secreted Gal-3 most likely binds to a ligand on the cell surface, which leads directly to the induction of the observed apoptotic response. Soluble Gal-3 interacts with over 30 lactosamine ligands on the cell surface, some of which are well characterized and include several extracellular matrix (ECM) components such as laminin, collagen IV, fibronectin vitronectin and elastin, as well as signalling receptors like CD4, CD66 and CD98, FcgRII, NCAM and Lamp-1 and 2. Additionally, it has been shown to bind to α1β1 and CD11b/ 18 integrins [9,30]. Clearly, further work is necessary to determine whether any of these known ligands can mediate the apoptotic effects of extracellular Gal-3 and may be expressed at higher levels in the tumor cells.

Examining the plasma levels of Gal-3 was especially necessary in the adjuvant setting, as patients had no residual tumor to follow post-surgery and only initial Gal-3 levels could be observed in the tissue. In the present study, only patients with a positive response to chemotherapy had a significant elevation in plasma Gal-3 levels. This may be due to the release of Gal-3 from circulating tumor cells that are unable to bind to partners normally present in the tumor microenvironment. Previously published results have shown higher levels of Gal-3 in cancer patients compared to normal controls [26-28], although these studies often involved patients who had received some form of cancer treatment, thus confounding the results. One study, Cheng et al. [28], examined Gal-3 in patients without any prior therapy but also limited analysis to serum Gal-3 levels prior to treatment. However, the Cheng et al. study measured initial Gal-3 levels consistent with the results of the present study. Regardless of such issues, Gal-3 levels in plasma have been suggested as a marker for metastatic potential [11,12,26-28]. Our results specifically examine responses to therapy and suggest that plasma levels of Gal-3 could be used to chart chemotherapy response when no visible tumor is present for disease monitoring.

The results further showed that a significant percentage of patients with elevated Gal-3 in response to adjuvant or neo-adjuvant chemotherapy also enjoyed longer disease-free survival over an 84-month follow up period, compared to patients with unchanged or decreased Gal-3 levels. Accordingly, we suggest that the increased plasma Gal-3 seen in previous studies may in fact be a response to cancer therapy, rather than a direct result of tumor cells themselves. Though these findings indicate clinical significance of Gal-3 as a marker of breast malignancy, *in vitro* research using chemo-sensitive and resistant breast cancer cell lines will be necessary to validate these results for further translation into clinical practice. Indeed, the study by Ilmer in 2016 performed several *in vitro* studies to elucidate the role of Gal-3 in Breast Cancer Stem Cells (BCSCs). That study found Gal-3 negative BCSCs to be highly tumorigenic and drug-resistant compared to the Gal-3 positive cells [8].

Finally, increased levels of stromal and plasma Gal-3 correlated with enhanced TUNEL staining post chemotherapy, indicating increased apoptosis in those tumors. Gal-3 has a well-established anti-apoptotic role in the cytoplasm, where it can antagonize the intrinsic cell death pathway via glycosylation-independent mechanisms [29]. Our results also implicate that soluble Gal-3 inhibits tumor cell growth through induction of apoptosis, partly by shuttling intracellular Gal-3 to the extracellular space. This could be due to p53 induced secretion of Gal-3 through exosome mediated pathways. Indeed, enhanced secretion of Gal-3 in response to p53 activation has been observed in a previous study [20]. We had examined p53 status of our study patients and had found a similar correlation between wt-p53 status and chemotherapy induced Gal-3 release but had a limited sample size (data not shown). Our present study had a small sample size due to the specific inclusion criteria and long follow up period. Still, even with the small cohort size a strong statistically significant relationship was seen between soluble Gal-3 levels and survival. Hence, further studies will be needed to confirm these results in a larger population.

A recent study has also shown strong apoptotic effect of a chimeric soluble Gal-3 designed to be secreted through addition of a classical secretion signal. In that study Lee et al. demonstrate that soluble Gal-3 kills aberrantly glycosylated tumor cells and antagonizes tumor growth through a novel integrin β1-dependent cell-extrinsic apoptotic pathway [30]. Similar post-translational modifications on β1integrin or other binding partners of Gal-3 found on the tumor cell surface may also be responsible for the pro apoptotic effect observed in our study. Based on our results and those of published peers we propose a mechanism where p53 activation in response to chemotherapy can induce release of Gal-3 leading to apoptotic response in effector as well as bystander cells containing aberrantly binding partners for Gal-3 (Fig. 6). Further studies would need to survey these modifications and binding partners in patient tissues to elucidate the pathways involved in the anticancer properties of soluble Gal-3. Future *in vitro* studies could also study the exact anti-tumor mechanism for soluble Gal-3 by examining the mobilization of Gal-3 in response to chemotherapy in cancer cell lines.

**Figure 6:**
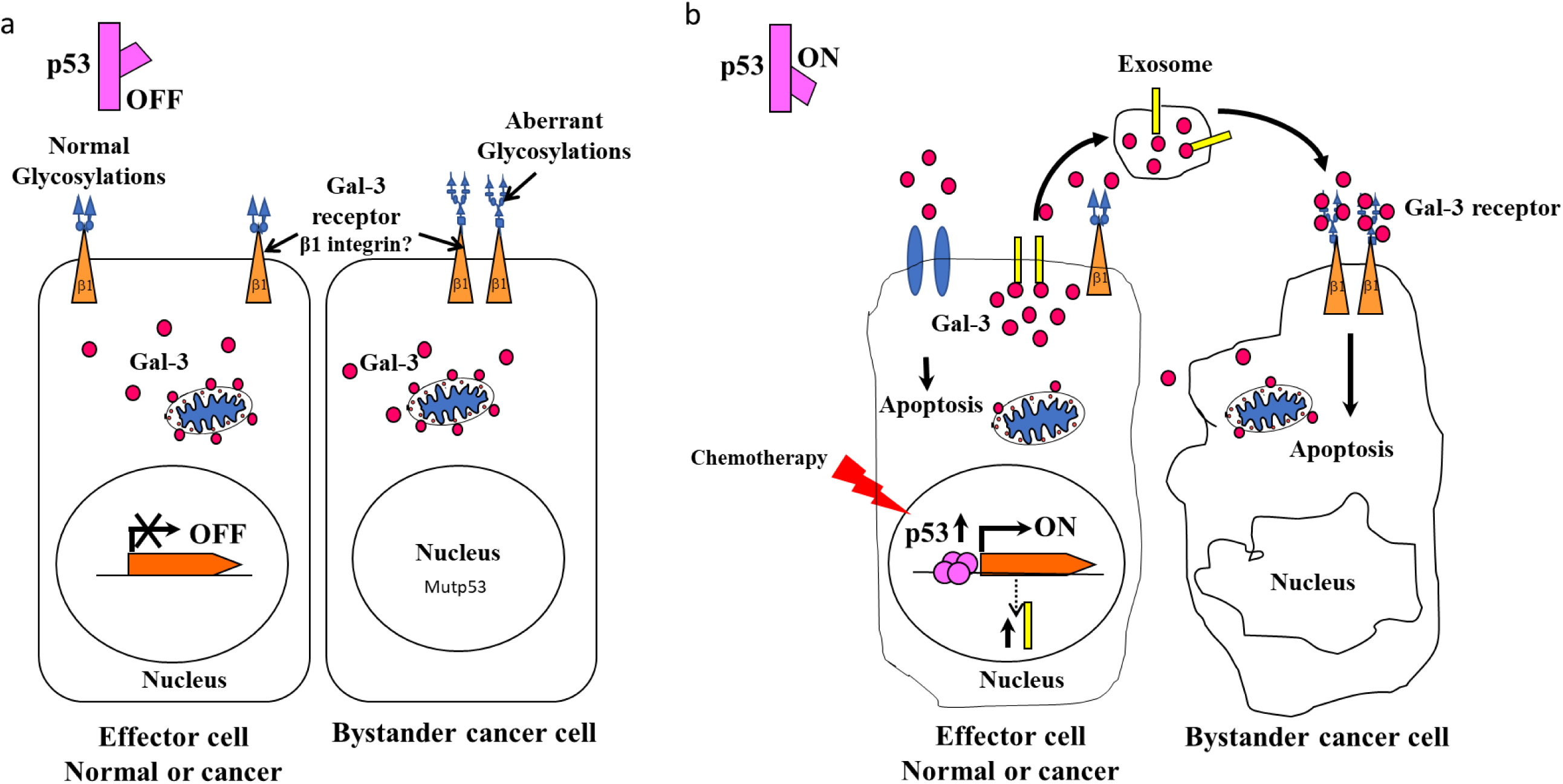
Proposed mechanism of action for chemotherapy induced extracellular Gal-3: a) Cells with inactive p53 have low extracellular, and high cytoplasmic levels of Gal-3. The cytoplasmic Gal-3 is anti-apoptotic through Bcl2 binding at the mitochondrial membrane. Many cancer cells have nonfunctional or partially functional p53 due to direct mutations or mutations in the pathway related to its activation. In addition, a major difference between normal and cancer cells is the presence of aberrantly-enhanced glycans on tumor cells. A recent study has found apoptotic response in cancer cells due to secreted Gal-3 binding to β1-integrin with aberrant N-glycosylations [30]. b) Activation of wt-p53 expression leads to secretion of Gal-3 through transcriptional upregulation of proteins like TSAP6 important for exosome formation [21]. This results in cytoplasmic Gal-3 secretion in the tumor environment with a concomitant decrease in intracellular levels. The secreted Gal-3 binds to β1-integrin with the abnormal glycosylations on same or bystander cell leading to apoptotic response. Lightning bolt represents chemotherapy induced activation of p53.

In summary, this study followed breast cancer patients who presented prior to any anticancer treatment in an attempt to clearly examine how surgery, chemotherapy and the disease itself alters secreted levels of Gal-3 in plasma and tumor stroma. Our results show a strong relationship between elevated Gal-3 levels in stroma and/or plasma of breast cancer patients and chemotherapy response. Therefore, monitoring extracellular and plasma Gal-3 levels can be used as a marker of chemotherapy efficacy. This is especially useful in adjuvant setting where no measurable tumor might be left to follow through typical imaging methods. We further provide the first evidence that enhanced levels of extracellular secreted Gal-3 in response to taxane therapy leads to better prognosis and longer disease-free survival. The results suggest the potential use of Gal-3 as an indicator for predicting long term patient prognosis.

## ACKNOWLEDGMENTS

We greatly appreciate those who volunteered to be a part of this study and donated their tissue and blood for our research. This work was supported with a research grant from Shaukat Khanum Memorial Cancer Hospital and Research Center to Fatima Khwaja Rehman (corresponding author). No conflict of interest is present for any of the authors.

## Ethical approval

All procedures performed in studies involving human participants were in accordance with the ethical standards of the institutional and/or national research committee and with the 1964 Helsinki declaration and its later amendments or comparable ethical standards.

## Informed consent

Informed consent was obtained from all individual participants included in the study.

## Author contributions

Project concept and experimental designs were developed by FKR, AS and JM. Patients were identified by NS; Patient recruitment, enrollment and sample procurement was done by AA; IHC analyses were analyzed and interpreted by SF, AS and AL; all experiments were performed by SF, AS and ALS and statistical analysis was done by OB.

