## Supplemental figures for "Elevated soluble Galectin-3 as a marker of chemotherapy efficacy in Breast cancer patients; a prospective study"

**Supplemental Material:**

**Supplemental Fig. 1:**

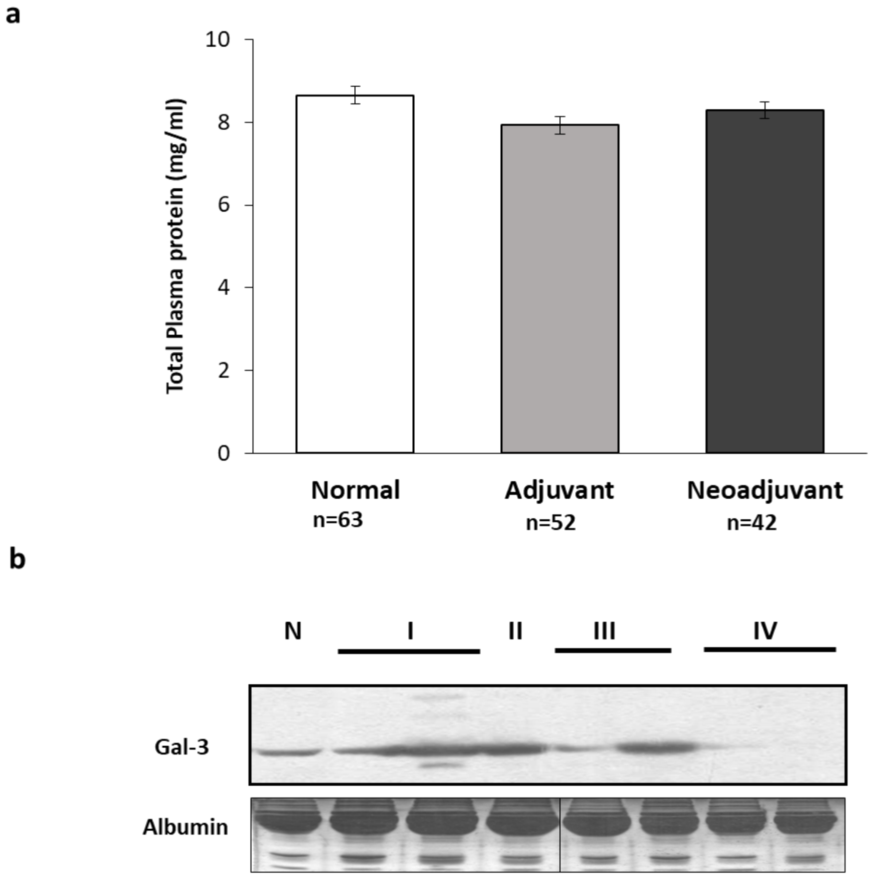


**Supplemental Fig. 1: Plasma levels of Galectin-3 decrease with increasing grade of tumor in breast cancer patients. a.** Total protein levels in plasma remain unchanged regardless of disease state in cancer patients prior to any treatment. **b.** A representative western blot showing relative Gal-3 expression in plasma from patients with varying grades of breast cancer and non-tumoral control. Albumin band on silver stained gel is used as a loading control.

**Supplemental Fig. 2**

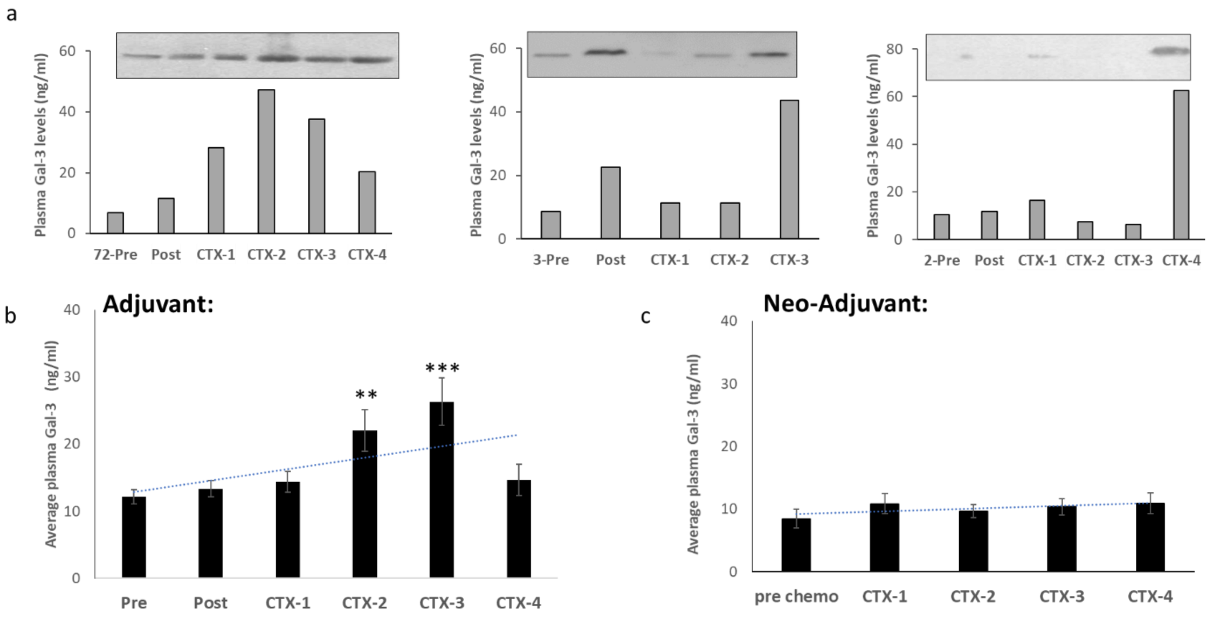


**Supplemental Fig. 2: Plasma Gal-3 levels increase in response to chemotherapy. a.** Representative ELISA quantification graphs and matched western blots confirming chemotherapy induced rise in plasma Gal-3 levels. Patient study number is listed at the beginning of each graph. Initial levels at the time of diagnosis (Pre), 2-weeks post surgical removal of tumor (Post) and after each chemotherapy cycle (CTX-1 to CTX-4) are shown. **b** Average plasma Gal-3 levels were calculated at the time of diagnosis (pre), 2 weeks post-surgical tumor removal (Post) and after each chemotherapy cycle (CTX-1-4) in patients receiving adjuvant chemotherapy (n=45). Notice significant increase in Gal-3 levels with each chemotherapy cycle in adjuvant patients (Pre:CTX2 p=0.008812; Pre:CTX3 p=0.000717 compared to initial (Pre) levels).

**Supplemental Fig. 3**

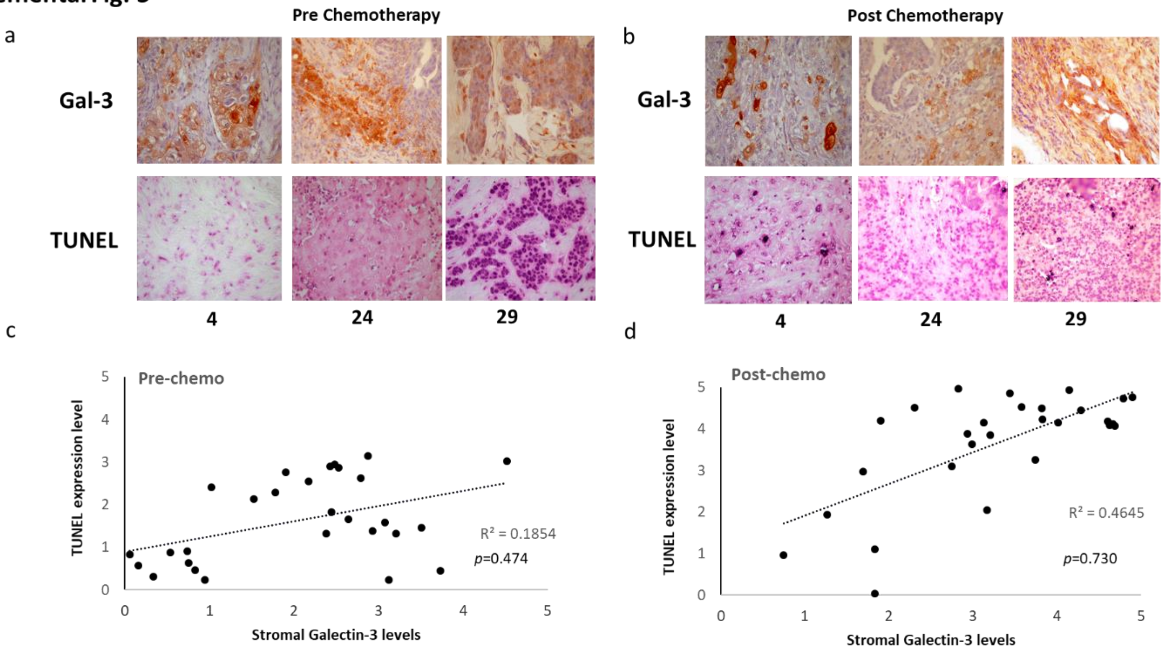


**Supplemental Fig. 3: Chemotherapy induced extracellular Gal-3 correlates with apoptosis in cancer cells. a.** TUNEL analysis of pre and post chemotherapy tumor tissue indicate positive relationship between stromal Gal-3 expression and apoptosis. Tissues with higher levels of stromal Gal-3 also show increase TUNEL reactivity. **b.** Correlation graph depicting the relationship between stromal Gal-3 expression and TUNEL reactivity. Pearson coefficient calculation was used to examine correlation. There was a strong positive correlation between stromal Gal-3 and TUNEL expressions post-chemotherapy (*p*=0.730). These results suggest an anti-tumor effect of soluble Gal-3.

**Supplemental Table I: Patient characteristics for breast cancer patients receiving Adjuvant chemotherapy after surgical removal of primary tumor.** Patient age, tumor size, type, grade and metastatic status are given. TRN=Triple negative; ER= Estrogen receptor; PR= Progesterone receptor; IDC=Intra-ductal carcinoma; DCIS= Ductal carcinoma in-situ; LCA= Lobular carcinoma; AC-T=Doxorubicin hydrochloride (Adriamycin) & Cyclophosphamide followed by Taxol; FAC/T=Fluorouracil, Doxorubicin, Cyclophosphamide followed by Taxol; CMF=Cyclophosphamide, Methotrexate, Fluorouracil.

| ADJUVANT Patient Info | | | | | | | | |
|---|---|---|---|---|---|---|---|---|
| **ID** | **Age** | **Tumor type** | **Grade** | **Ln status** | **ER** | **PR** | **Her2Neu** | **Chemo type** |
| **1** | 41 | IDC | III | + | - | - | 2+ | AC/ T |
| **2** | 29 | IDC | III | + | - | - | - | AC/T |
| **3** | 39 | IDC | III | - | - | + | - | CMF/T |
| **6** | 34 | IDC | II | + | - | - | 3+ | AC/ T |
| **7** | 43 | IDC | II | + | + | + | - | CMF/ T |
| **8** | 53 | IDC | II/ III | + | - | - | + | FAC/T |
| **9** | 64 | IDC | II | + | + | + | - | FAC/ T |
| **10** | 43 | IDC | II | - | + | + | 2+ | CMF/T |
| **46** | 63 | IDC | II | + | + | - | - | FAC/T |
| **47** | 29 | IDC | II | + | - | - | - | AC/T |
| **48** | 37 | IDC | III | + | - | + | - | AC/T |
| **49** | 43 | IDC | II | + | - | - | - | FAC/T |
| **50** | 59 | IDC | III | N/A | + | + | + | AC/T |
| **51** | 54 | IDC | III | + | - | - | - | AC/T |
| **52** | 32 | IDC | II | + | + | + | 2+ | AC/T |
| **53** | 36 | IDC | II | + | - | - | + | AC/T |
| **54** | 44 | IDC | III | - | + | + | - | FAC |
| **55** | 44 | IDC | II | + | + | + | 2+ | AC/T |
| **56** | 28 | IDC | III | + | - | - | - | AC/T |
| **57** | 38 | IDC | III | + | - | - | - | FAC |
| **58** | 59 | IDC | III | - | N/A | N/A | N/A | AC/T |
| **59** | 51 | IDC | III | + | + | - | 2+ | FAC/T |
| **60** | 54 | IDC | III | + | + | - | - | FAC |
| **68** | 39 | IDC | III | + | + | - | 3+ | AC/T |
| **71** | 37 | IDC | II | + | 75% + | 90% + | | Taxol |
| **72** | 49 | IDC | II | + | + | + | - | AC/T |
| **73** | 40 | IDC | III | + | - | - | - | AC/T |
| **74** | 44 | IDC | III | + | + | - | - | FAC/T |
| **75** | 42 | IDC | II | + | - | + | 1+ | AC/T |
| **76** | 39 | Sarcoma | III | + | N/A | N/A | N/A | AC/T |
| **77** | 34 | IDC | III | - | - | - | - | AC/T |
| **78** | 56 | ILC | II | - | N/A | N/A | N/A | AC/T |
| **79** | 54 | IDC | III | - | N/A | N/A | N/A | AC/T |
| **80** | 51 | IDC | III | - | N/A | N/A | N/A | AC/T |
| **81** | 57 | IDC | III | - | - | - | - | AC/T |
| **82** | 46 | IDC | II | + | + | + | - | AC/T |
| **83** | 56 | ILC | II | + | N/A | N/A | N/A | AC/T |
| **84** | 59 | IDC | II | - | + | + | - | AC/T |

| | | | | | | | | |
|---|---|---|---|---|---|---|---|---|
| **85** | 64 | IDC | II | + | N/A | N/A | N/A | FAC/T |
| **86** | 59 | IDC | III | + | + | - | 3+ | FAC/T |
| **87** | 34 | IDC | III | - | + | + | 2+ | AC/T |
| **88** | 39 | IDC | III | - | + | + | - | FAC |
| **89** | 47 | ILC | IV | + | - | 30% + | - | AC/T |
| **90** | 45 | IDC | III | + | <10% | <10% | - | AC/T |
| **91** | 46 | IDC | II | N/A | N/A | N/A | N/A | AC.T |
| **101** | 55 | IDC | II | - | - | - | 3+ | AC/T |
| **102** | 44 | IDC | III | + | <10% | <10% | - | AC/T |
| **103** | 33 | IDC | III | + | - | - | 2+ | AC/T |
| **104** | 47 | IDC | II | + | 20-40% + | 10-15% + | - | AC/T |
| **105** | 65 | IDC/ ILC | II | + | 80% + | 50% + | - | AC/T |
| **106** | 55 | IDC | II | - | 10% + | - | 2+ | FAC/T |
| **107** | 56 | IDC | II | + | 10%+ | 10%+ | 2+ | AC/T |

TRN=Triple negative; ER= Estrogen receptor; PR= Progesterone receptor; IDC=Intra-ductal carcinoma; DCIS= Ductal carcinoma in-situ; LCA= Lobular carcinoma; AC-T=Doxorubicin hydrochloride (Adriamycin) & Cyclophosphamide followed by Taxol; FAC/T=Fluorouracil, Doxorubicin, Cyclophosphamide followed by Taxol; CMF=Cyclophosphamide, Methotrexate, Fluorouracil.

**Supplemental Table II: Patient characteristics for breast cancer patients receiving Neo-adjuvant chemotherapy followed by surgery.** Patient age, tumor size, type, grade and metastatic status are given. TRN=Triple negative; ER= Estrogen receptor; PR= Progesterone receptor; IDC=Intra-ductal carcinoma; DCIS= Ductal carcinoma in-situ; LCA= Lobular carcinoma. AC-T=Doxorubicin hydrochloride (Adriamycin) & Cyclophosphamide followed by Taxol; FAC/T=Fluorouracil, Doxorubicin, Cyclophosphamide followed by Taxol; CMF=Cyclophosphamide, Methotrexate, Fluorouracil.

| Neo-adjuvant Patient Info | | | | | | | | |
|---|---|---|---|---|---|---|---|---|
| **ID** | **Age** | **Tumor type** | **Grade** | **Ln status** | **ER** | **PR** | **Her2Neu** | **Chemo type** |
| **4** | 58 | IDC | III | - | - | - | - | FAC/T |
| **5** | 35 | IDC | III | - | + | + | - | FAC/T |
| **21** | 26 | IDC | N/A | - | - | - | 2+ | AC/T |
| **22** | 48 | Metastatic Ca | N/A | + | - | - | - | AC/T |
| **23** | 49 | IDC | III | + | - | - | 3+ | AC/T |
| **24** | 41 | IDC | III | + | - | - | 3+ | FAC/T |
| **25** | 39 | IDC | III | + | 30% | 20% | 3+ | FAC/T |
| **26** | 39 | ILC | II | + | - | - | 3+ | FAC/T |
| **27** | 52 | IDC | III | + | - | - | - | AC/T |
| **28** | 40 | IDC | III | + | - | - | - | AC/T |
| **29** | 48 | IDC | II | - | 60-70 | 70-80 | - | AC/T |
| **30** | 39 | IDC | III | - | + | + | - | AC/T |
| **67** | 59 | IDC | II | + | 80 | 90 | - | AC/T |
| **69** | 47 | IDC | I | - | 35-40 | - | - | Carboplatin/ T |
| **70** | 62 | IDC | II | + | 50 % | 10% | - | AC/T |
| **94** | 44 | IDC | II | - | - | - | 3+ | FAC/T |
| **98** | 54 | ILC | II | + | + | + | - | FAC/ T |
| **99** | 44 | IDC | III | + | - | - | - | AC/T |
| **92** | 41 | LCIS | III | - | + | - | - | AC/T |
| **93** | 47 | IDC/ ILC | II | - | 5% | 5% | - | AC/T |
| **95** | 43 | IDC/ ILC | II | - | 5% | 5% | - | AC/T |
| **96** | 48 | IDC | III | + | - | - | - | AC/T |
| **97** | 52 | IDC | III | - | - | - | - | AC/T |
| **100** | 20 | IDC | III | + | - | - | 2+ | FAC/ T |
| **108** | 54 | IDC | IV | N/A | N/A | N/A | N/A | FAC/ 5FU/ T |
| **109** | 39 | IDC | II | - | 60% | 30% | N/A | AC/ T |
| **110** | 42 | IDC | III | + | 50% | - | - | AC/ T/ Doc |
| **111** | 55 | IDC | III | + | N/A | N/A | N/A | FAC/T |
| **112** | 50 | IDC | III | + | - | - | - | AC/ T |
| **113** | 39 | IDC | II | - | - | - | + | AC/ T |
| **114** | 30 | IDC | III | + | - | - | - | Carboplatin/ AC/T |
| **115** | 45 | IDC | II | N/A | - | - | 3+ | FAC/T |
| **116** | 34 | IDC | II | N/A | - | - | - | AC/T |
| **117** | 63 | IDC | III | N/A | N/A | N/A | N/A | |
| **118** | 49 | IDC | II | - | - | - | - | AC/ T |
| **119** | 45 | IDC | III | + | - | - | 3+ | AC/T |
| TRN=Triple negative; ER= Estrogen receptor; PR= Progesterone receptor; IDC=Intra-ductal carcinoma; DCIS= Ductal carcinoma in-situ; LCA= Lobular carcinoma; AC-T=Doxorubicin hydrochloride (Adriamycin) & Cyclophosphamide followed by Taxol; FAC/T=Fluorouracil, Doxorubicin, Cyclophosphamide followed by Taxol; CMF=Cyclophosphamide, Methotrexate, Fluorouracil. | | | | | | | | |
